# Tree Species as Biomonitors of Air Pollution around a Scrap Metal Recycling Factory in Southwest Nigeria: Implications for Greenbelt Development

**DOI:** 10.1101/2024.02.06.579208

**Authors:** D. G. Olanipon, F. K. Ayandeyi, A. E. Enochoghene, O. A. Eludoyin, B. A. Adanikin, O. O. Awotoye

## Abstract

Trees are biomonitors and sinks for air pollutants but better sinking ability comes from trees with high tolerance for air pollution. Consequently, this study investigated the Air Pollution Tolerance Index (APTI) and Anticipated Performance Index (API) of six dominant tree species around a scrap metal recycling factory in Ile-Ife, Southwest Nigeria. Biochemical and physiological parameters such as the relative water content, total chlorophyll, leaf extract pH and ascorbic acid content of the leaves of the selected tree species were determined and used to compute the APTI. The biological and socio-economic characters of each tree species were equally examined to determine the API. The APTI of the selected tree species during the dry season was in the N. *laevis* (11.8) > *A*. *boonei* (11.2) > *S. siamea* (11.0) > *B. micrantha* (10.8) *> T. orientalis* (10.6) *> T. grandis* (9.6). According to the API grading, *N*. *laevis* and *A*. *boonei* were classified as “good” (62.5% each) tree species for greenbelt development for both dry and wet seasons, while *T*. *orientalis* was also classified as a “good” (62.5% each) tree species for greenbelt development for the wet season only. Native tree species such as *N*. *laevis*, *A*. *boonei* and *T*. *orientalis* exhibited better tolerance to gaseous pollutants and are recommended for biomonitoring environmental health and greenbelt establishment.

## Introduction

Increased industrial activities have culminated in increased air pollution in globally (Karine *et al*., 2011; Ahmed *et al*., 2019; Amegah, 2020). Urbanization, transportation and establishment of industries that process raw materials into finished products are major sources of gaseous pollutants Developmental activities such as these are necessary because of the services they provide. Used metals and plastics which are dumped indiscriminately lead to accumulation of pollutants such as heavy metals in the soil, air and water environment.

A major means of reducing waste accumulation and conserving the environment is via recycling. Recycling is the production of a useable product from a waste material. This is a necessary process to manage certain wastes (Shi *et al*., 2021) such as plastics, iron and other metals in both rural and urban settings. Over the years, scrap metal wastes have been recycled via the iron smelting process.

Emission of pollutants (particulate matter, CO_2_, SO_2_, iron oxides, alumina, silica, alkali oxides and heavy metals in oxidized gaseous forms) are associated with iron smelting processes during the recycling process (Brook *et al*., 2004; Pope & Dockery, 2006; Zhang *et al*., 2009; Tai *et al*., 2010; Symanski *et al*., 2020). The iron smelting process is a recycling process that great health concern to fauna and people living within the vicinity of metal recycling complex (Owoade *et al*., 2009; Solademi and Thompson, 2020) just as it was observed in the study area. These categories of pollutants are air-based and travel over long distances in the air before being deposited (Enyoh *et al*., 2019).

Oxides of nitrogen and sulphur (SO_X_ and NO_X_) in the air dissolve in rain water to form sulphuric and nitric acid respectively, causing acid rain (Sivakumaran 2015; Ali, 2021). The acid rain leads to corrosion of buildings, monuments and statues, causes soil acidity, and lead to soil degradation (Liu *et al*., 2019). Other air pollutants like carbon dioxide (CO_2_) and methane (CH_4_) trap the heat radiated from the earth and raise the earth’s temperature of a phenomenon referred to as global warming (Venkataramanan & Smitha, 2011; Rajak, 2021). Short and long term exposures to air pollutants have also been linked with premature mortality and reduced life expectancy in humans (Kampa & Castanas, 2008; Dedoussi *et al*., 2020).

The use of vegetation to purify the atmosphere of air pollutants is fast gaining acceptance, due of their enormous leaf area for impingement, absorption and accumulation of air pollutants in plant tissues (Escobedo *et al*., 2008; Lui & Ding, 2008; Das, 2010; Xing & Brimblecombe, 2019; Diener & Mudu, 2021; Araminienė *et al*., 2023). Trees are major air pollutants sinks (Oksanen & Kontunen-Soppela, 2021), and their presence around indusrtial areas such as the metal recycling factory could reduce the impact of air pollution from such facility on humans and animals. Identifying tree species suitable for green belt is a function of assessing the ability of tree species that can tolerate extreme pollution levels without obvious effects on their metabolism, physiology and growth. Although, all trees are affected by air pollution but the degree of the impact vary from species to species (Singh *et al*., 2021). Some of these effects leaf injury, stomata damage, premature senescence, decreased photosynthetic activities, lowered membrane permeability and reduced growth and yield in sensitive plant species (Tiwari *et al*., 2006).

Trees act as air pollution sinks but the better performance comes from the pollution tolerant species (Miria and Khan, 2013; Roy *et al*., 2020). The response of the plant’s physiological and biochemical characters to air pollution can be understood by analyzing the factors determining resistance and susceptibility (Govindaraju *et al*., 2012; Salimi & Dadkhah Aghdash, 2022). The objectives of the this research are to (i) determine the physiological and biochemical parameters of tree species to determine their Air Pollution Tolerance Indices (ii) to investigate the socioeconomic value of the trees species to determine their Anticipated Performance Indices. This study therefore compared the response and tolerance of some selected tree species to gaseous and particulate pollutants within the vicinity of the scrap metal recycling factory to identify appropriates species for greenbelt development.

## Materials and methods

### Study Area

The study was carried out within the vicinity of an iron and steel recycling factory (latitude 7° 29’ 30’’ N and 7° 30’ 0’’ N and longitude 4° 28’ 30’’ E and 4° 29’ 30’’ E) (Figure 1), along Ife-Ibadan expressway about 3 km from Ife-Ibadan-Ilesha roundabout and about 5 km from the Obafemi Awolowo University main campus, Ile-Ife, Osun state, Nigeria.

**Figure 1:**
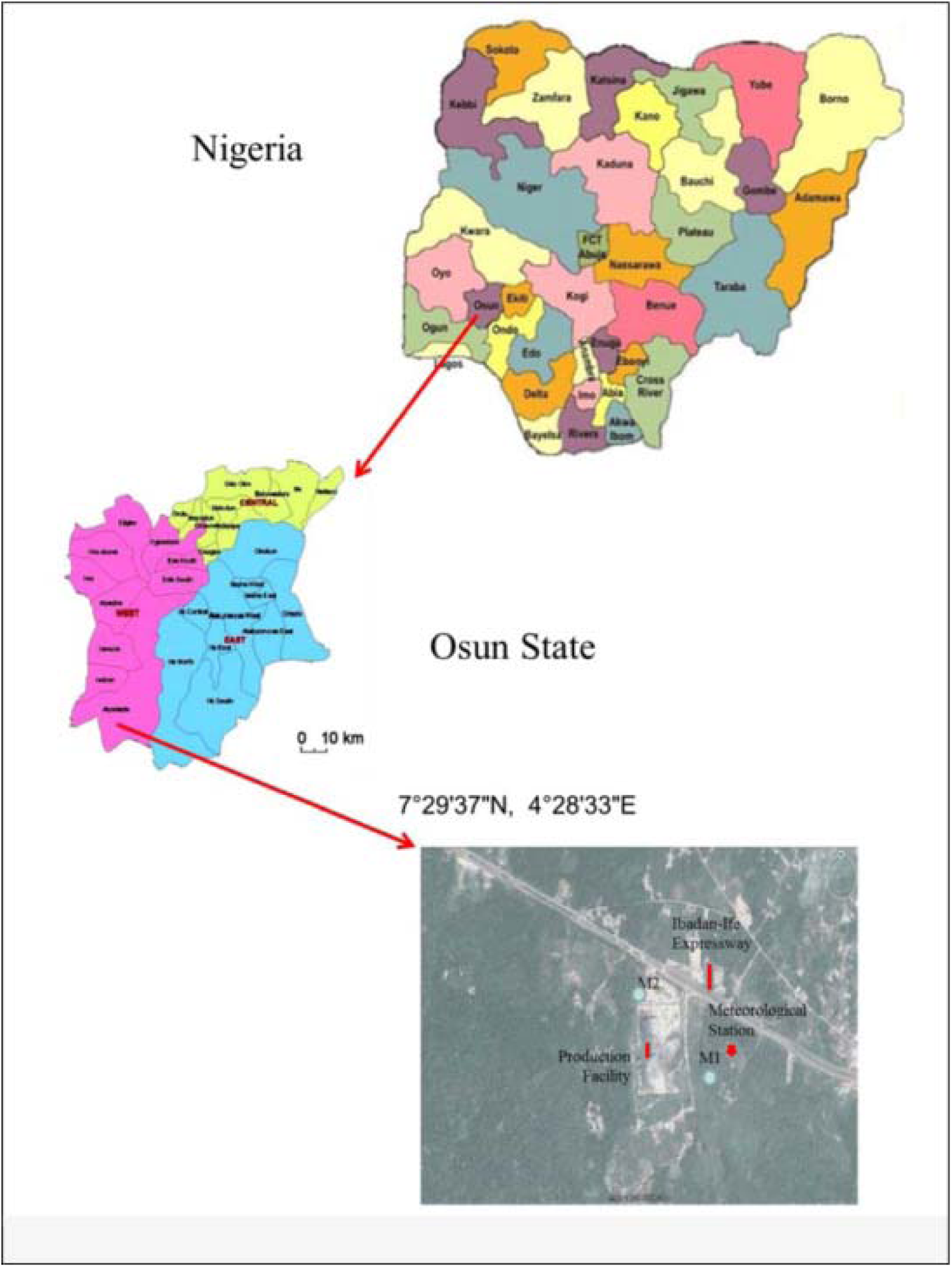
Map of Nigeria showing the study area (Source: Owoade et al., 2015)

### Sampling

Sampling plots were established along line transects at increasing distance from the Iron and Steel Recycling Factory, Ile-Ife. The transects were located in the four cardinal directions of North (N), East (E), South (S), and West (W) of the factory. The plots were established starting from the zero meters, (factory site) up to three hundred and fifty meters in each of the four directions. Dominant tree species whose height were about 1.3 m and above, with a girth size of 10 cm and above (which indicates that the stand had become fully matured) were purposively selected and used for the study.

### Leaf Sample Collection

Matured fresh leaf samples were collected in triplicates from each tree species that were selected for the study. The leaf samples were collected between the hours of 7:00 am to 11:00 am for six days at intervals two weeks in both dry and wet seasons. The leaf samples were transferred immediately to the laboratory for determination of its fresh weight. Thereafter, biochemical analysis of parameters such as the relative water content, total chlorophyll, leaf extract pH, and ascorbic acid content of the leaves of the selected tree species were determined.

### Relative Water Content

The relative water content was determined using the method described by Tripathi *et al*. (2009). Fresh weight was obtained by weighing the fresh leaves on a weighing balance (Model Scout Pro SPU 2000). The leaves were then immersed in water overnight, blotted dry with a blotting paper and then weighed to determine the turgid weight. Thereafter, the leaves were dried overnight in an oven at 70 ^0^C and weighed to obtain the dry weight.

The relative water content (% RWC) was determined using the formula:

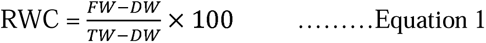

Where:

FW is the fresh weight (g).

DW is the dry weight (g) of the turgid leaves after oven drying at 70^0^C.

TW is the turgid weight (in grams) after immersion in water overnight.

### Leaf Extract pH

The leaf extract pH was determined with a glass electrode pH meter (Model PHS – 3B). Distilled water (50 ml) was added to 5 grams of grinded leaf sample. The pH of the obtained suspension was measured with a pH meter after calibrating the pH meter with buffer solution of pH 4 and 9 (Adamsab & Kousar, 2011).

### Ascorbic Acid (AA) Content

Ascorbic acid content which is usually expressed in mg/g was measured using spectrophotometric method (Begum and Harikrishna, 2010). One gram of fresh foliage was added to a test-tube containing 4 ml oxalic acid-Ethylene-diamine-tetra-acetic acid (0.05 M oxalic acid, 0.2 M EDTA) extracting solution, 1 ml of orthophosphoric acid, 1ml sulphuric acid (5%), 2 ml of ammonium molybdate and 3 ml of distilled water. The solution was then left for 15 minutes and absorbance was read at 760 nm with a spectrophotometer (Bio Quest UV spectrophotometer Cecil 2000 series). The concentration of ascorbic acid in the samples was then extrapolated from a standard ascorbic acid curve.

### Total Chlorophyll (TCh)

The total chlorophyll content in the leaves was estimated following the method of Liu & Ding, (2008). Three grams of fresh leaves was blended and the chlorophyll was extracted with 10ml of 80% acetone and left for 15 minutes. The liquid portion was decanted into another test tube and centrifuged at 2500 rpm for 3 minutes. The supernatant was then collected and the absorbance read at 645 nm (D_645_) and 663 nm (D_663_) using a spectrophotometer (Bio Quest UV spectrophotometer Cecil 2000 series). The optical density of the total chlorophyll (CT) is the sum of chlorophyll a (D_645_) density and chlorophyll b (D_663_) density. Thus,

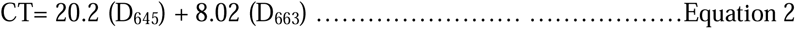

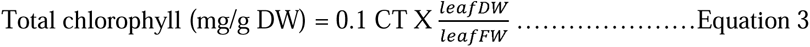

Where: 0.1 = constant

CT = optical density of the total chlorophyll

### Determination of Air Pollution Tolerance Index (APTI) of Plants

The air pollution tolerance index (APTI) proposed by Singh & Rao, (1983) to assess the tolerance of plants to air pollution was used. The air pollution tolerance index (APTI) was calculated from values obtained during the biochemical analysis, using the formula:

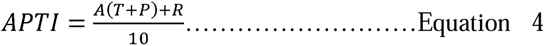

Where: A = ascorbic acid (mg/g)

T = total chlorophyll (mg/g)

P = pH of the leaf extract

R = relative water content of leaf (%)

10 = constant

Based on the resultant APTI values gotten from the biochemical parameters analysed in the leaves of the tree species, the plants were grouped into three categories as shown in Table 2

**Table 1.** Location of the Selected Tree Species at the Experimental site.

| Scientific name | Family | North | South | East | West |
| --- | --- | --- | --- | --- | --- |
| *Newbouldia laevis* | Bignoniaceae | + | - | - | - |
| *Trema orientalis* | Ulmaceae | - | + | - | - |
| *Tectona grandis* | Verbanaceae | - | + | - | - |
| *Bridelia micrantha* | Phyllanthaceae | - | - | + | - |
| *Alstonia boonei* | Apocynaceae | - | - | - | + |
| *Senna siamea* | Fabaceae | - | - | - | + |

+ = present
- = absent

**Table 2:** Determination of Tolerance Level of Tree Species.

| APTI | VALUE | RESPONSE |
| --- | --- | --- |
| A | <11 | Sensitive |
| B | 12 – 16 | Intermediate |
| C | 17 and above | Tolerant |

(Source: Padmavathi *et al.*, 2013)

### Determination of Anticipated Performance Index (API)

The anticipated performance index (API) was calculated for different tree species by combining the resultant APTI values with some relevant biological and socio-economic characters of the plant (plant habit, type of plant, canopy structure, laminar structure and economic value). Based on these characters, different grades (+ or –) were allotted to plants and allotted points were scaled to a percentage system to produce categories of assessment as shown in Appendix 1. The criteria that were used for calculating the API of different plant species are shown in Table 3.

**Table 3.** Mean Values of Air Pollution Tolerance Index of the Selected Tree Species during Dry and Wet Seasons.

| Tree species | Dry season | Response | Wet season | Response |
| --- | --- | --- | --- | --- |
| *Newbouldia laevis* | 11.8<sup>a</sup> ± 0.5 | Intermediate | 11.9<sup>a</sup> ± 0.8 | Intermediate |
| *Trema orientalis* | 10.6<sup>b</sup> ± 0.8 | Sensitive | 13.3<sup>a</sup> ± 0.4 | Intermediate |
| *Tectona grandis* | 9.6<sup>b</sup> ± 0.9 | Sensitive | 11.7<sup>a</sup> ± 0.5 | Intermediate |
| *Bridelia micrantha* | 10.8<sup>b</sup> ± 0.4 | Sensitive | 12.9<sup>a</sup> ± 0.3 | Intermediate |
| *Alstonia boonei* | 11.2<sup>a</sup> ± 0.7 | Intermediate | 10.9<sup>b</sup> ± 0.2 | Sensitive |
| *Senna siamea* | 11.0<sup>a</sup> ± 0.5 | Sensitive | 12.9<sup>a</sup> ± 0.3 | Intermediate |

Mean (± Standard Error) for each variable with the same alphabet across the columns are not significantly different (p<0.05).

Based on this grading system, a tree species can only secure a maximum of 16 positive points, and allotted points were scaled to a percentage system to produce categories of assessment.

### Statistical Analysis

The statistical analyses of data collected during the experiment were carried out using software SPSS 20.0. The data were analysed with Graphical tools and Correlation and results were considered significant at 95% confidence interval.

## Results

For the biochemical parameters carried out in the leaves of the various tree species, it was observed that the pH of the leaf extracts were within the acidic range both in the dry and wet season, with *Bridelia micrantha* being more acidic in both seasons (Figure 2). However, the leaves of the tree species exhibited lower pH in the dry season. In Figure 3, significant difference occurred only in the ascorbic acid content of *Trema orientalis* and *Bridelia micrantha* during the dry and wet seasons. It was observed that the ascorbic acid content of *Alstonia boonei when compared in the two seasons,* had a higher content in the dry season than the wet season relative to other tree species. *Newbouldia laevis* had the highest total chlorophyll among the tree species and seasonal variation in the total chlorophyll content of the studied tree species had no significant effect (Figure 4). In Figure 5, the RWC of the trees was higher in the wet season except in *Newbouldia laevis* and *Alstonia boonei*.

**Figure 2:**
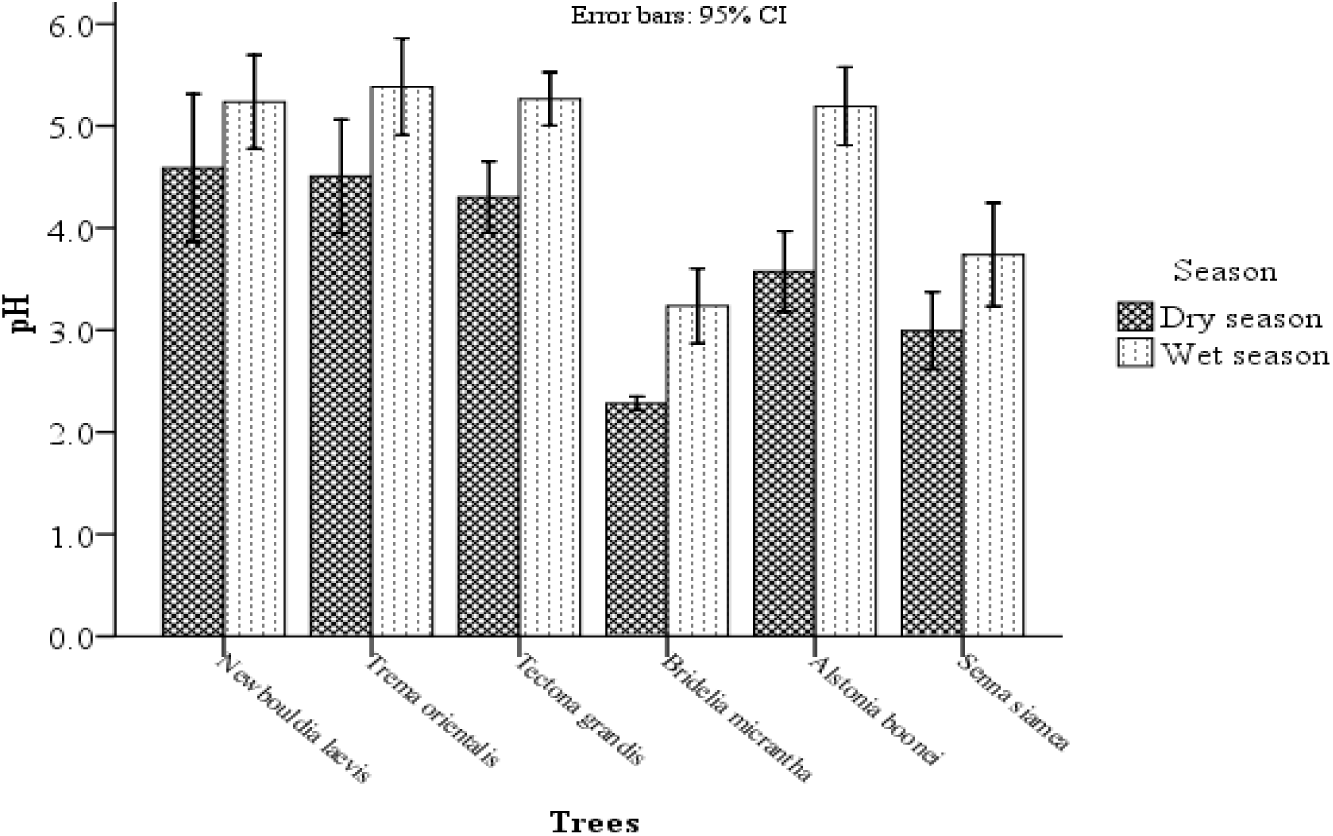
Seasonal variation in pH of the tree species

**Figure 3.**
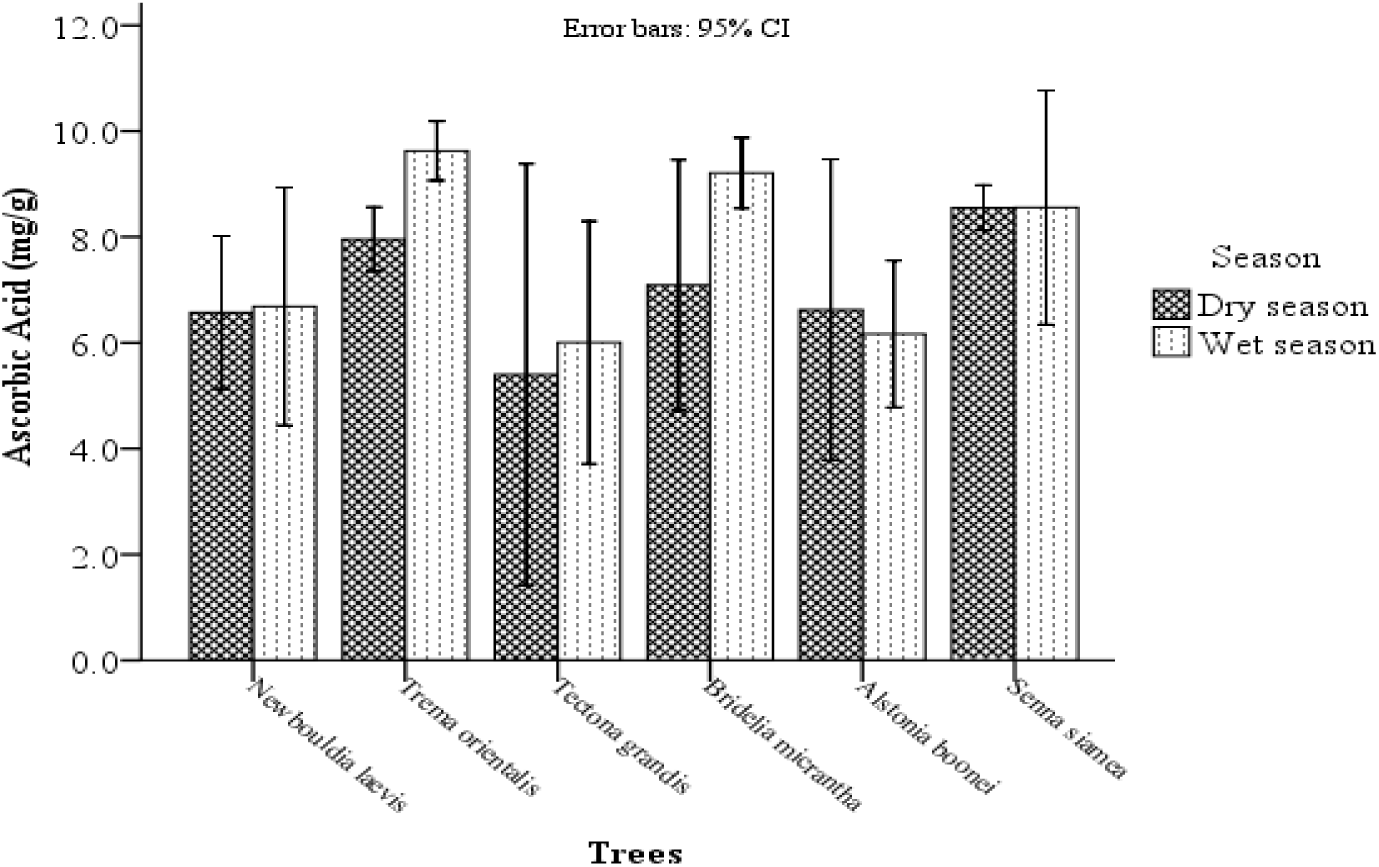
Seasonal variation of ascorbic acid in the tree species

**Figure 4.**
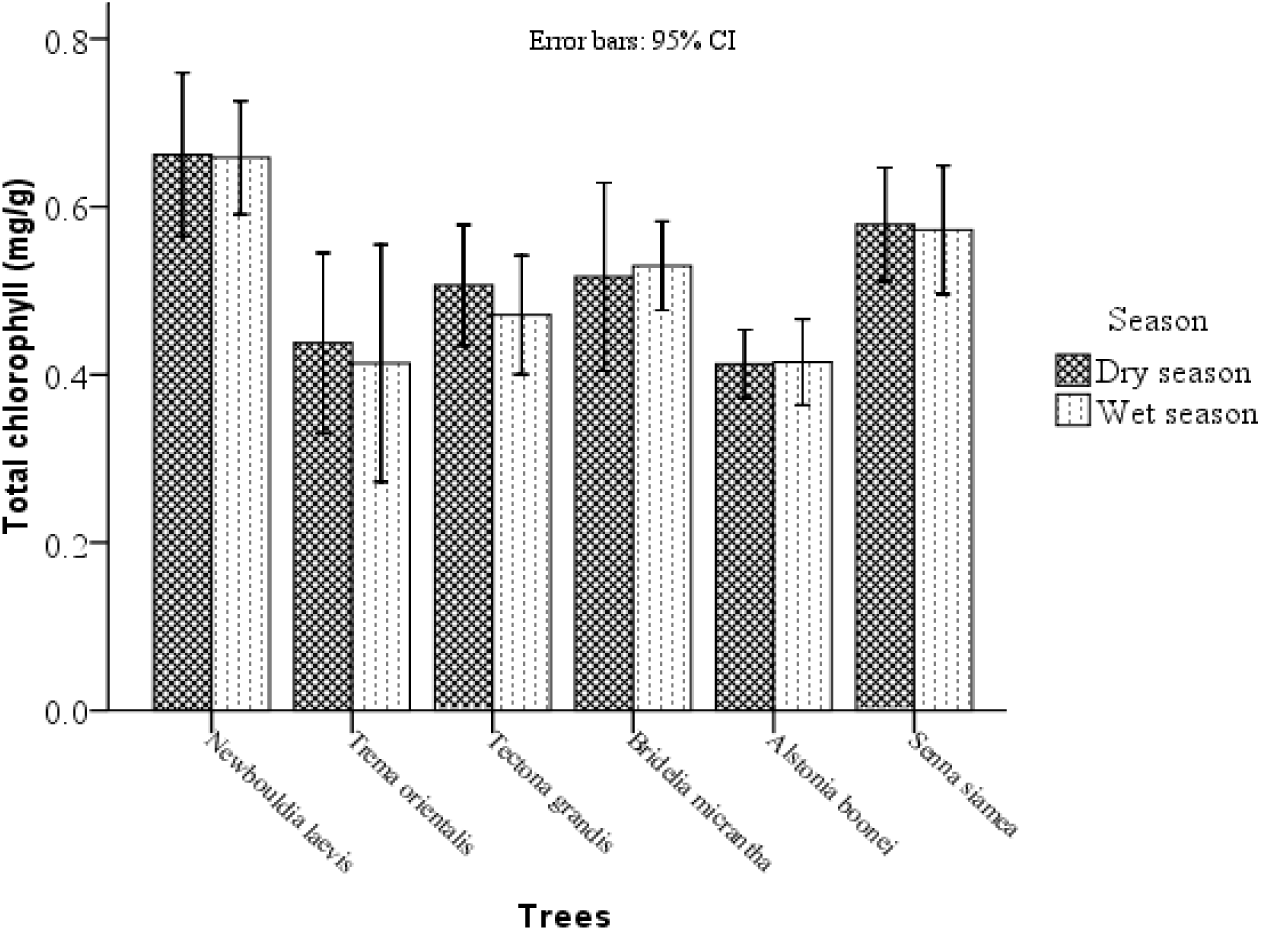
Seasonal variation of total chlorophyll in the tree species

**Figure 5.**
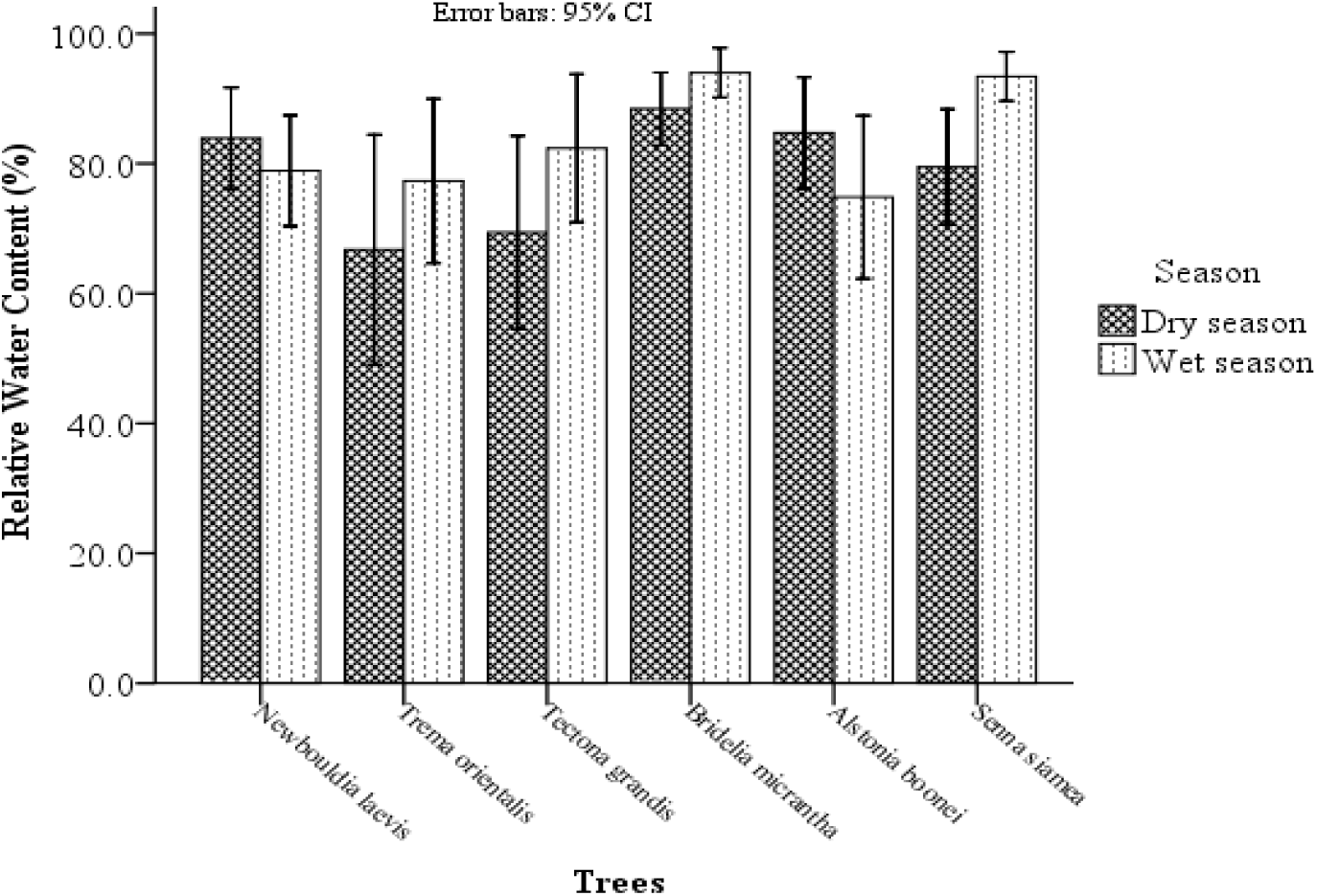
Seasonal variation of relative water in the tree species

### Sensitivity and Tolerance Levels of Selected Trees

As shown in Table 3, at both seasons, the APTI of *N*. *laevis* was adjudged to be intermediate. The APTI also indicated that four of the six trees under study namely: *Trema*, *Tectona Bridelia* and *Senna* were sensitive during the dry season and intermediate in the wet season. *Alstonia* was the only tree species considered to be intermediate during the dry season and sensitive during the wet season.

### Relationship between Biochemical Parameters and Air Pollution Tolerance Index (APTI) in the Dry and Wet Seasons

The relationship among the biochemical parameters and APTI in the two seasons studied are shown in Tables 4 and 5. A strong relationship was observed between the ascorbic acid content and APTI in the dry (r = 0.652) and wet (r = 0.808) seasons. Also, relative water and APTI had a strong relationship in the dry (0.656) and wet (0.665) seasons.

**Table 4.** Correlation between the four biochemical parameters and APTI as observed in the dry season.

|  | pH | Ascorbic Acid | Total Chlorophyll | Relative Water Content | APTI |
| --- | --- | --- | --- | --- | --- |
| pH | 1 | -0.220 | 0.055 | -0.327 | 0.045 |
| Ascorbic Acid |  | 1 | -0.213 | 0.057 | 0.652* |
| Total Chlorophyll |  |  | 1 | 0.059 | -0.112 |
| Relative Water Content |  |  |  | 1 | 0.656* |
| APTI |  |  |  |  | 1 |

**Table 5.** Correlation between the four biochemical parameters and APTI as observed in the wet season.

|  | pH | Ascorbic<br>Acid | Total<br>Chlorophyll | Relative Water<br>Content | APTI |
| --- | --- | --- | --- | --- | --- |
| pH | 1 | -0.374 | -0.102 | -0.653* | -0.322 |
| Ascorbic Acid |  | 1 | 0.065 | 0.261 | 0.808* |
| Total<br>Chlorophyll |  |  | 1 | 0.157 | 0.062 |
| Relative Water<br>Content |  |  |  | 1 | 0.665* |
| APTI |  |  |  |  | 1 |

indicate that there is a strong relationship at p<0.01

### Greenbelt Assessment

In order to obtain the Anticipated Performance Index (API) of each tree species data was obtained on the various biological and socioeconomic values of each species plant habitat, canopy structure, type of plant, laminar structure and APTI. Following a grading scale (Appendix 2) proposed by Govindaraju *et al*. (2012), six plant species were evaluated according to a grading pattern to determine their anticipated performance index.

The results of the assessment of the selected tree species for greenbelt development classified *Newbouldia leavis* and *Alstonia boonei* as “good” (62.5% each) performers for greenbelt development during the dry season, while *Trema orientalis* and *Tectona grandis* were classified as “moderate” with total score of about 56.3% each. Also, *Senna siamea* and *Bridelia micrantha* having total grading percentage score of 50% and 43.8% respectively were classified as “poor” performers in the dry season (Table 6).

**Table 6:**
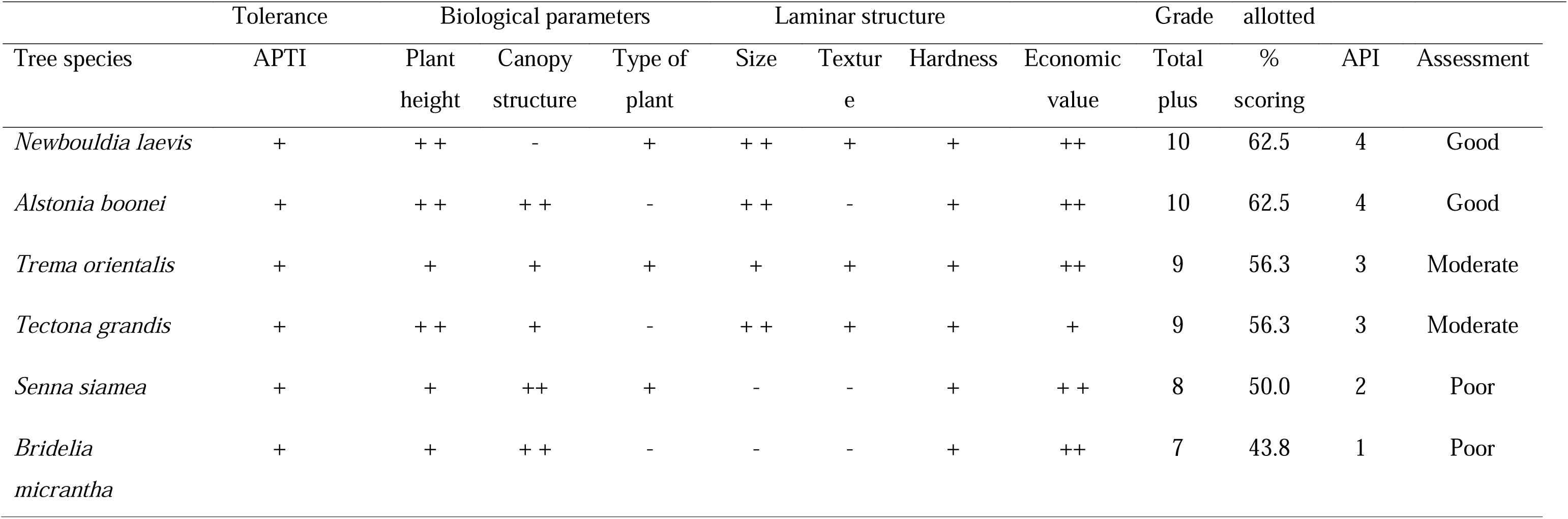
Identification of Tree Species for Greenbelt Development (Dry Season Observation)

In the wet season, *Newbouldia laevis, Alstonia* sp. and *Trema orientalis* were classified as “good” (62.5% each) plant species for greenbelt development, while *Tectona grandis* and *Senna siamea* were classified as “moderate” with total score of about 56.3% each. However, *Bridelia micrantha* having a total grading percentage score of 43.8% is classified as a “poor” performer (Table 7). By interpretation, *Newbouldia laevis*, *Alstonia boonei*, *Trema orientalis* and *Tectona grandis* are identified as appropriate tree species for greenbelt assessment (Tables 6 and 7).

**Table 7.** Identification of Tree Species for Green Belt Development (Wet Season Observation)

| Tree species | Tolerance | Biological parameters |  |  | Laminar structure |  |  |  | Grade | allotted |  |  |
| --- | --- | --- | --- | --- | --- | --- | --- | --- | --- | --- | --- | --- |
|  | APTI | Plant height | Canopy structure | Type of plant | Size | Texture | Hardness | Economic value | Total plus | % scoring | API | Assessment |
| *Newbouldia laevis* | + | ++ | - | + | ++ | + | + | ++ | 10 | 62.5 | 4 | Good |
| *Alstonia boonei* | + | ++ | ++ | - | ++ | - | + | ++ | 10 | 62.5 | 4 | Good |
| *Trema orientalis* | ++ | + | + | + | + | + | + | ++ | 10 | 62.5 | 4 | Good |
| *Tectona grandis* | + | ++ | + | - | ++ | + | + | + | 9 | 56.3 | 3 | Moderate |
| *Senna siamea* | ++ | + | ++ | + | - | - | + | ++ | 9 | 56.3 | 3 | Moderate |
| *Bridelia micrantha* | ++ | + | ++ | - | - | - | + | ++ | 8 | 50.0 | 1 | Poor |

## Discussion

Plant species such as trees, shrubs and herbs act as bioindicators (detectors) and biomonitors of pollution in the environment (Ogunrotimi et al., 2017; Diener & Mudu, 2021).). The tolerance level of any plant to pollution is indicated by the effect of the pollutants on the plant’s physiological and biochemical parameters such as photosynthetic activity as revealed by the chlorophyll content, acidity, water and mineral levels (Adak & Kour, 2021). A plant species that exhibits tolerance to pollution is able to continue its growth and physiological activities, while a sensitive plant species may become susceptible to changes caused by the pollutants and experience cell death, and eventual senescence

The pH of leaves of the tree species investigated was generally acidic both in the dry and wet season. This may be attributed to the level of SO_2_ and NO_x_ present in the ambient air, has previous studies have shown that these gases can change in the pH of the leaves towards acidity (Swami *et al.,* 2004; Punit and Rai, 2021). The acidic nature of the leaves also show that gaseous pollutants, such as S0_2_and NO_X_ undergo diffusion, and react with cellular water to form acid radicals.(Mondal *et al*. 2011; Kousar *et al*. 2014; Adak and Kour, 2021). The acidic state of the leaves correlates well with a sensitive APTI, and has been shown to lower the rate of photosynthesis in such plants(Yan-Ju and Hui, 2008; Thakar and Mishra, 2010; Karmakar et al., 2021). This is in line with this study, where all the tree species had a relatively low leaf extract pH. A similar report by Tsega & Devi (2014) also confirmed this, as all plant species exposed to vehicular emissions had low pH values.

The slightly high concentration of Ascorbic acid (AA) concentration in both the dry and wet season is similar to the findings Kousar *et al*. (2014) and Rai (2020), where it was recorded that plants resistant to pollutants contained high amount of ascorbic acid, while plants sensitive to pollutants contained rather low ascorbic acid. Findings in the current study showed *Tectona grandis* and *Alstonia boonei* which are classified as sensitive plants according to the APTI grading, with a lower mean AA concentration than the other tree species studied. This showed that plants with a high ascorbic acid content exhibit hriger tolerance to pollutants when exposed to them. (Kousar *et al.,* 2014; Ter *et al*., 2020). Hence, sufficient amount of ascorbic acid in plant tissues and organs helps plants resist adverse environmental conditions. (Lima *et al*., 2000; Karmakar *et al*., 2021).

The present study revealed the reduced amount of chlorophyll in all the plant species and this can be attributed to exposure of the leaf area to oxides of sulphur, nitrogen and Ozone, this further reduces the sites for photosynthetic activity (Agrawal *et al*., 2003; Asif *et al*., 2021). Acid radicals are formed in the leaf matrix as these oxides react with cellular water and thus affecting formation of chlorophyll molecules. Hence, such plant species may become intolerant to pollution, experience cell death and eventually die-off (Sharma & Saxena, 2022). In this study, the chlorophyll content of the plant species did not vary significant as the seasons changed, as the low leaf extract pH also causes reduction in the chlorophyll content.

Relative Water Content (RWC) in this study was high, averaging above 70%. Plants exposed to air pollution stress are still able maintain their physiological balance and survive with optimum water content in their tissues (Singh *et al*., 2020). In addition, RWC in plants determines protoplasmic permeability of cells, thereby controlling water loss and amount of nutrient dissolved nutrients; too low water content may result in leaf senescence (Tsega and Devi-Prasad, 2014). Hence, high water content in plant tissues presents an advantage for proper functioning of biological processes in polluted environments (Meerabai *et al*., 2012; Chaudhary *et al*., 2021).

The APTI value of each tree species was computed using the four biochemical parameters in plant leaves viz; pH, ascorbic acid, total chlorophyll value and relative water content, this values also helps in can predicting the air quality of an environment. Plants having higher APTI value are classified as tolerant to air pollution while plants with low APTI value are classified as less tolerant (Bhadauria *et al*., 2022). Karmakar & Padhy (2019) opined that a plant species may be sensitive to air pollution in one geographical location while it may be tolerant in another area, this can be attributed to prevailing weather and climate conditions of the area under study. The present study clearly shows that plant species respond differently to air pollution, in additions that no plant species have the highest value for all the four biochemical parameters, and that each parameter plays a unique role in the establishing the susceptibility or tolerance of plants. Hence, the purpose of combining the four biochemical parameters in determining the air pollution tolerance index of each species.

APTI and ascorbic acid and relative water content exhibited a strong positive correlation in the study, this was also observed by Mondal *et al*. (2011) and Bui *et al*. (2021) who observed that a high positive correlation exist between APTI and ascorbic acid and relative water content in their assessment of woody tree species to air pollution. This correlation points to the fact that ascorbic acid and relative water content are the highly crucial factors in determining the tolerance to plant species to air pollution.

Comparison of APTI values for different plants revealed the sensitivity and tolerance of the tree species. It was reported that trees with relatively low APTI value exhibit sensitivity to air pollutants while those high APTI value exhibit tolerance to air pollution (Sadia *et al*., 2019). Among the six tree species studied, *Newbouldia laevis* and *Alstonia boonei* were intermediate (moderately tolerant to air pollution) and other tree species were sensitive in the dry season, while *Alstonia boonei* was the only sensitive tree species and others were intermediate in the wet season. In addition, *Senna siamea* had an APTI value which is approximately similar to the value obtained by Prajapathi and Tripathi (2008) for *Senna siamea* (14.5) found in an urban area, Varanisi city in India. Based on APTI value tree species classified as “sensitive” can be used as bio-indicators, while those classified as “tolerant” tree species can be planted as greenbelts to serve as sinks for air pollutants.

Findings from the study reveal that *Newbouldia laevis*, *Alstonia boonei* and *Trema orientalis* can be anticipated to be a “good” performer, while *Tectona grandis* and *Senna siamea* are predicted to be “moderate” greenbelt performers and *Bridelia micrantha* is classified as “poor” greenbelt performer. Similar results were obtained by Anake *et al*. (2019) who sampled three tree species from an industrial environment in Southeast Nigeria. Several researchers from Nigeria and other countries have identified tree species such as *Newbouldia laevis, Tectona grandis, Terminalia catappa Polyalthia longifolia, Mangifera indica, Ficus benghalensis, Saraca asoka* and several others as excellent and good greenbelt performers (Ogunrotimi et al., 2017; Karmakar & Padhy; 2019; Karmakar *et al*. 2021).

## Conclusion

Therdetermination of anticipated performance index of tree species is highly useful in the selection of appropriate species for development of greenbelt development. This study therefore established that *Newbouldia laevis*, *Trema orientalis* and *Alstonia boonei* are ideal species for biomonitoring, greenbelt development especially in polluted areas such as industries, so as to sequester gaseous pollutant, clean the air and reduce greenhouse effect. Tree species such as *Trema orientalis* and *Tectona grandis* that were classified as moderate performers can be recommended in afforestation efforts and plantation forestry due their economic value and their ability to serve as air filters for biomonitoring in polluted environments.

## Notes

### Competing Interest Statement

The authors have declared no competing interest.

