## Supplementary information for "Tree Species as Biomonitors of Air Pollution around a Scrap Metal Recycling Factory in Southwest Nigeria: Implications for Greenbelt Development"

Supplementary figure 1

Gradation of plant species on the basis of air pollution tolerance index (APTI) and other biological and socio-economic characters.

| Grading character | | Pattern of assessment | Grading allotted |
| --- | --- | --- | --- |
| 1. Tolerance | APTI | 9.0 – 12.0  12.1 – 15.0  15.1 – 18.0  18.1 – 21.0  21.1 – 24.0 | +  + +  + + +  + + + +  + + + + + |
| 2. Biological and socioeconomic | Plant height  Canopy structure  Type of plant | Small  Medium  Large  Sparse/irregular/globular  Spreading crown/open/semi-dense  Spreading dense  Deciduous  Evergreen | –  +  + +  –  +  + +  –  + |
| 3. Laminar structure (Leaf) | Size  Texture  Hardness  Economic value | Small  Medium  Large  Smooth  Coriaceous  Delineate  Hardy  < 3 uses  3 – 5 uses  Above 5 uses | –  +  + +  –  +  –  +  –  +  + + |

Prajapati and Tripathi (2008); Govindaraju *et al*. (2012).

Supplementary figure 2

Grading Scale of Anticipated performance index (API) of plant species

| Grade | Scores (%) | Assessment category |
| --- | --- | --- |
| 0 | < 30 | Not recommended |
| 1 | 31 – 40 | Very poor |
| 2 | 41 – 50 | Poor |
| 3 | 51 – 60 | Moderate |
| 4 | 61 – 70 | Good |
| 5 | 71 – 80 | Very good |
| 6 | 81 – 90 | Excellent |
| 7 | 91 – 100 | Best |

Source: Prajapati and Tripathi (2008); Govindaraju *et al*., (2012).
